# Precision at Every Scale: Efficiency in AI-Driven *De Novo* Antibody Design

**DOI:** 10.1101/2025.11.21.689414

**Authors:** Hyeonjin Cha, Kyesoo Cho, Jeonghyeon Gu, Daehyeon Gwak, Seung Woo Ham, Mirim Hong, Sehan Kim, Suyeon Kim, Sohee Kwon, Changsoo Lee, Da Kyoung Lee, Danyeong Lee, Dohoon Lee, Jungsub Lim, Jinsung Noh, Soyeon Oh, Eunhwi Park, Seongchan Park, Taeyong Park, Eunwoo Ryu, Seongok Ryu, Deok Hyang Sa, Chaok Seok, Jiho Sim, Moo Young Song, Jonghun Won, Hyeonuk Woo, Jinsol Yang

**Affiliations:** Galux Inc, Seoul 08758, Republic of Korea

## Abstract

The precise *de novo* design of antibodies remains a therapeutic challenge. The AI platform, GaluxDesign, was evaluated in a high-efficiency Precision-Scale Workflow by synthesizing and testing only 50 full-length IgG candidates per epitope across eight distinct epitopes from six therapeutic targets. This campaign yielded a 10.5% binder rate (estimated EC_50_ *<* 100 nM), identifying target-specific binders for seven of eight epitopes, with multiple candidates exhibiting sub-nanomolar to single-digit nanomolar dissociation constants (K_d_). We further assessed the same workflow on nine shared benchmark targets selected for external comparison, where GaluxDesign identified target-specific binders for eight of nine targets, demonstrating strong target-level performance relative to previously reported *de novo* antibody design approaches. Together, these results establish a high-efficiency, precision-scale workflow for generating novel, high-affinity therapeutic antibodies.

## 1 Introduction

The precise *de novo* design of antibodies, which engage targets through six variable loops, remains a critical challenge in therapeutic molecule discovery. Unlike conventional methods relying on existing immune repertoires, AI-driven *in silico* design offers the capacity to generate novel binders with precisely tailored molecular properties.

The practical realization of this potential requires a design methodology of validated versatility and fidelity. The GaluxDesign platform [1, 2] addresses this by leveraging two distinct, complementary discovery modalities: a Precision-Scale Workflow (rapid, high-fidelity validation using minimal candidates) and a Library-Scale Workflow (comprehensive functional exploration of sequence space).

Our prior work successfully validated the Library-Scale approach (10^6^ candidates), demonstrating the platform’s extensive capacity for generating diverse, functional antibody repertoires [1]. This report validates the complementary component: the high precision and efficiency of the GaluxDesign platform. We demonstrate that our *in silico* design framework enables the Precision-Scale Workflow, confirming this approach as a rapid, resource-efficient method for identifying therapeutic-grade antibodies from a minimal candidate pool, thereby completing the robust demonstration of our platform’s versatile capabilities.

## 2 Results and Discussion

### 2.1 Workflow Setup

To validate the high efficiency of the GaluxDesign platform [1], we established a rigorous, small-scale validation workflow. The workflow started with the selection of eight distinct epitopes across six therapeutically relevant target proteins for *de novo* design. These targets included Programmed Death-Ligand 1 (PD-L1), Human Epidermal Growth Factor Receptor 2 (HER2), Activin type 2B receptor (ACVR2B), and Activin receptor-like kinase 7 (ALK7). Notably, two distinct epitopes were selected for Interleukin-11 (IL-11): one targeting the IL-11RA binding site (IL-11_R*α*_) and the other targeting the GP130 binding site (IL-11_GP130_), chosen for their potential for pH selectivity. Similarly, two epitopes were selected for Frizzled class receptor 7 (Fzd7): a novel epitope (Fzd7_site1_) chosen to confer subtype specificity among the Frizzled family, and another epitope (Fzd7_site2_) situated near a hydrophobic patch utilized by known binders (PDB ID: 6NE4 [3], 6O3A [4]), which had not been previously explored in our *de novo* design campaigns.

From the computationally generated pool, we selected the top 50 *de novo* candidates per epitope (400 total designs) for experimental synthesis and validation (Figure 1). This minimal pool size was chosen to stringently assess the predictive fidelity and ranking power of the *in silico* modules. Furthermore, all designs were expressed directly as full-length Immunoglobulin G (IgG) molecules, bypassing intermediate formats to provide a rigorous and therapeutically relevant validation.

**Figure 1.**
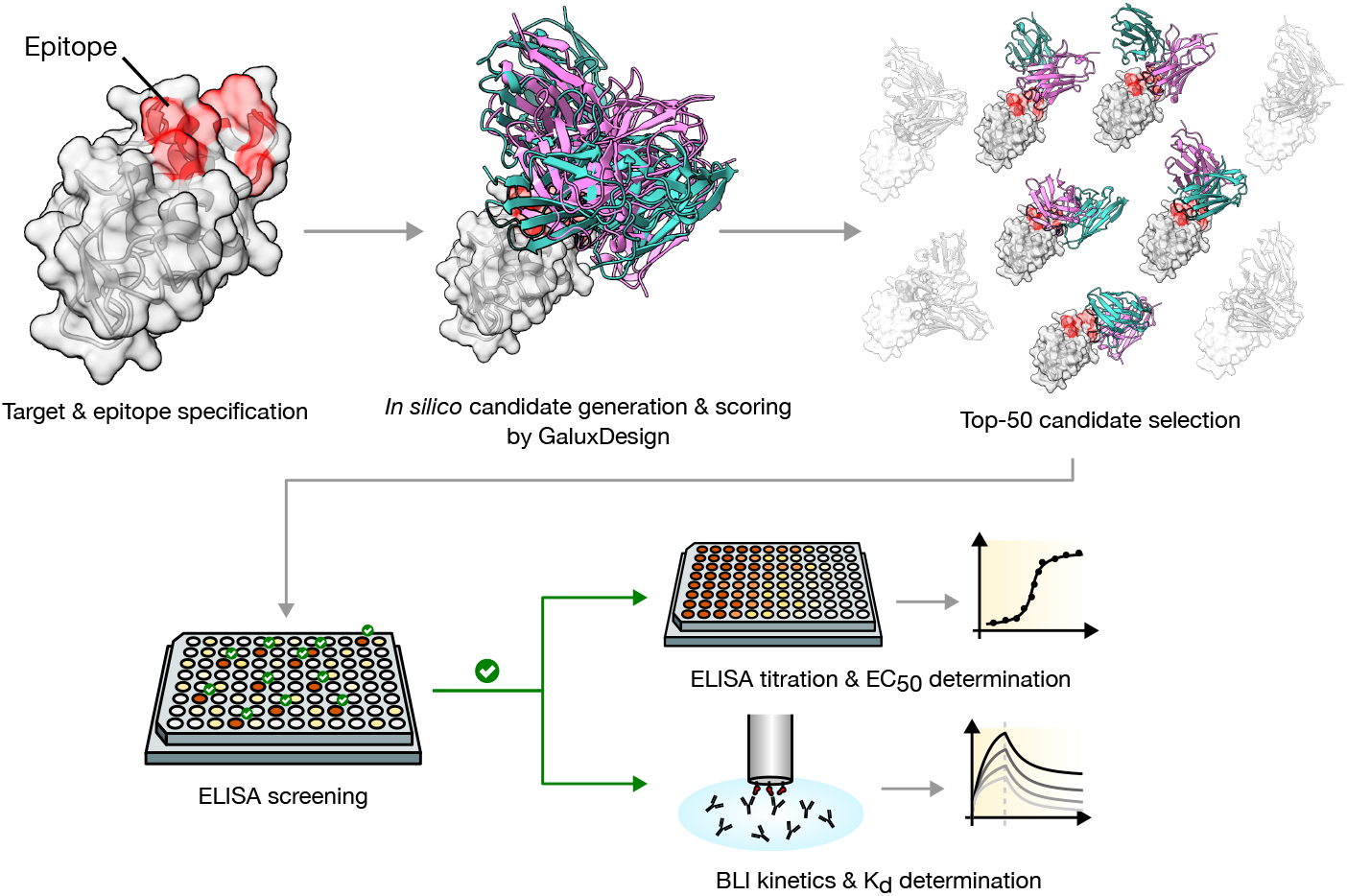
Schematic of the High-Efficiency *De Novo* Antibody Discovery Workflow. The process begins with target and epitope specification, followed by *in silico* candidate generation using GaluxDesign platform. A computational scoring function then selects the top 50 candidates per epitope for experimental validation. These candidates are synthesized as full-length IgG molecules for initial ELISA screening. Confirmed binders subsequently undergo detailed biophysical characterization, including EC_50_ determination and BLI K_d_ measurement.

Seven of the eight epitopes were previously validated targets in our Library-Scale (10^6^) screening workflow. Their inclusion facilitates a direct comparison, demonstrating that the platform’s advanced high-precision scoring can efficiently identify candidates from a minimal pool. The eighth epitope (Fzd7_site2_) was included to confirm the platform’s generalizability to newly-defined target sites. The design strategy relied solely on the target protein structure (including the predicted model for ALK7) and designated epitope residues.

### 2.2 Performance Validation: High Success Rates in the Precision-Scale Workflow

The precision-scale workflow yielded exceptional performance metrics, decisively demonstrating the high efficiency and predictive power of the GaluxDesign platform. From the 400 total designs tested, we achieved an overall binders(relaxed) rate of 31.5% and, most notably, an overall binders(strict) rate of 10.5% (Figure 2). The binders(relaxed) criteria were set to align with similar metrics reported in other *in silico de novo* design works [5, 6, 7, 8], while the more stringent binders(strict) criteria, defined as estimated EC_50_ *<* 100nM, were intended to impose a higher standard for qualifying candidates as potential hits. For detailed information on binder definition and classification, see Subsection 4.3 and Supplementary Figure S8.

**Figure 2.**
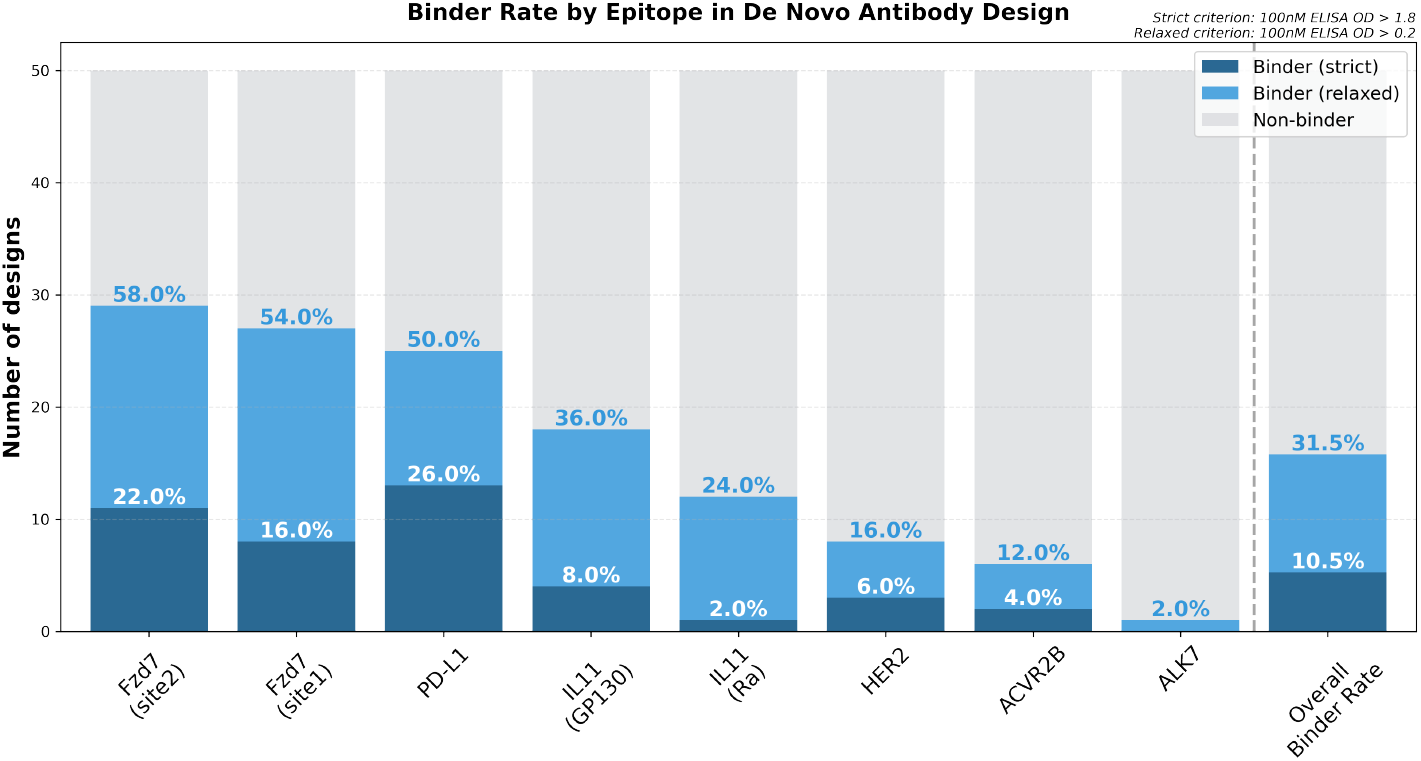
Binder Rate by Epitope in *De Novo* Antibody Design. Results from a precision-scale validation campaign testing 50 *de novo* IgG design candidates per epitope. Bars represent the number of designs classified as binders(strict) (100 nM ELISA OD *>* 1.8, dark blue), binders(relaxed) (100 nM ELISA OD *>* 0.2, light blue), or Non-binders (grey). The overall binder rates across all eight epitopes were 10.5% (strict) and 31.5% (relaxed) (far-right bar).

The platform’s reliability was further confirmed by its success across the targets. Strict binders were successfully identified for seven of the eight targeted epitopes (87.5% success rate). This included the newly-defined Fzd7_site2_, which achieved a 22.0% binders(strict) rate. For ALK7, a challenging target lacking an experimental structure, the workflow still generated binders(relaxed) (2.0%) against the computationally predicted structure, confirming measurable and specific binding.

### 2.3 Sequence Novelty and Structural Diversity of Designed Binders

We assessed the novelty and diversity of our discovered binders at both the sequence and structural levels (Figure 3). The analysis confirmed a high degree of sequence novelty; a substantial fraction of the identified binders exhibits a maximum Complementarity-Determining Region (CDR) sequence identity of *<* 65% compared to any antibody structure in the PDB (SAbDab)[9] (Figure 3A). This indicates that GaluxDesign successfully generates candidates in sequence spaces outside of known antibody repertoires.

**Figure 3.**
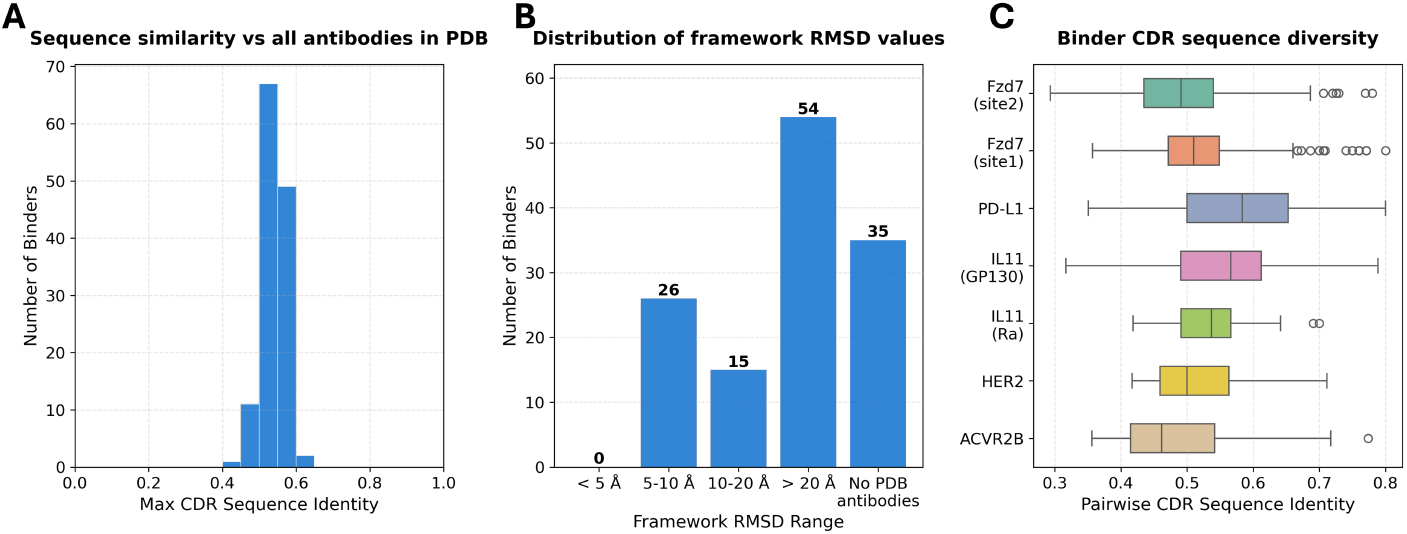
Novelty and Diversity of *in silico* Designed Binders. (A) CDR sequence identity comparison of the designed binders against all antibodies in the PDB. (B) Distribution of Framework C-*α* RMSD values for the binders compared to the nearest known PDB complex for the same target. (C) Pairwise CDR sequence identity distribution within the binders(relaxed) pools for each epitope, demonstrating high diversity among the generated binders.

The novelty of binding pose was confirmed by aligning the target protein from our predicted complex to the nearest PDB complex. The resulting distribution of Framework C-*α* Root Mean Square Deviation (RMSD) values indicates that a substantial fraction of the binders adopts binding orientations that are structurally distinct from previously characterized binding modes (Figure 3B). Furthermore, analysis of the pairwise CDR sequence identity among binders confirmed significant sequence diversity within the hit pools for each epitope (Figure 3C).

### 2.4 *In Vitro* Characterization: High Affinity and Epitope Fidelity

A subset of high-affinity binders underwent comprehensive *in vitro* characterization (Table 1). The detailed kinetic analysis confirmed that the platform consistently produces high-affinity molecules, with affinities ranging from sub-nanomolar to single-digit nanomolar dissociation constants (K_d_) confirmed for five of the eight targeted epitopes. Sub-nanomolar affinities were confirmed for key targets: PD-L1 (K_d_ = 0.1 nM), Fzd7 (K_d_ = 0.28 nM), ACVR2B (K_d_ = 0.46 nM), and HER2 (K_d_ = 0.87 nM), with a single-digit nanomolar binder also validated for IL-11 (K_d_ = 4.2 nM).

**Table 1.**
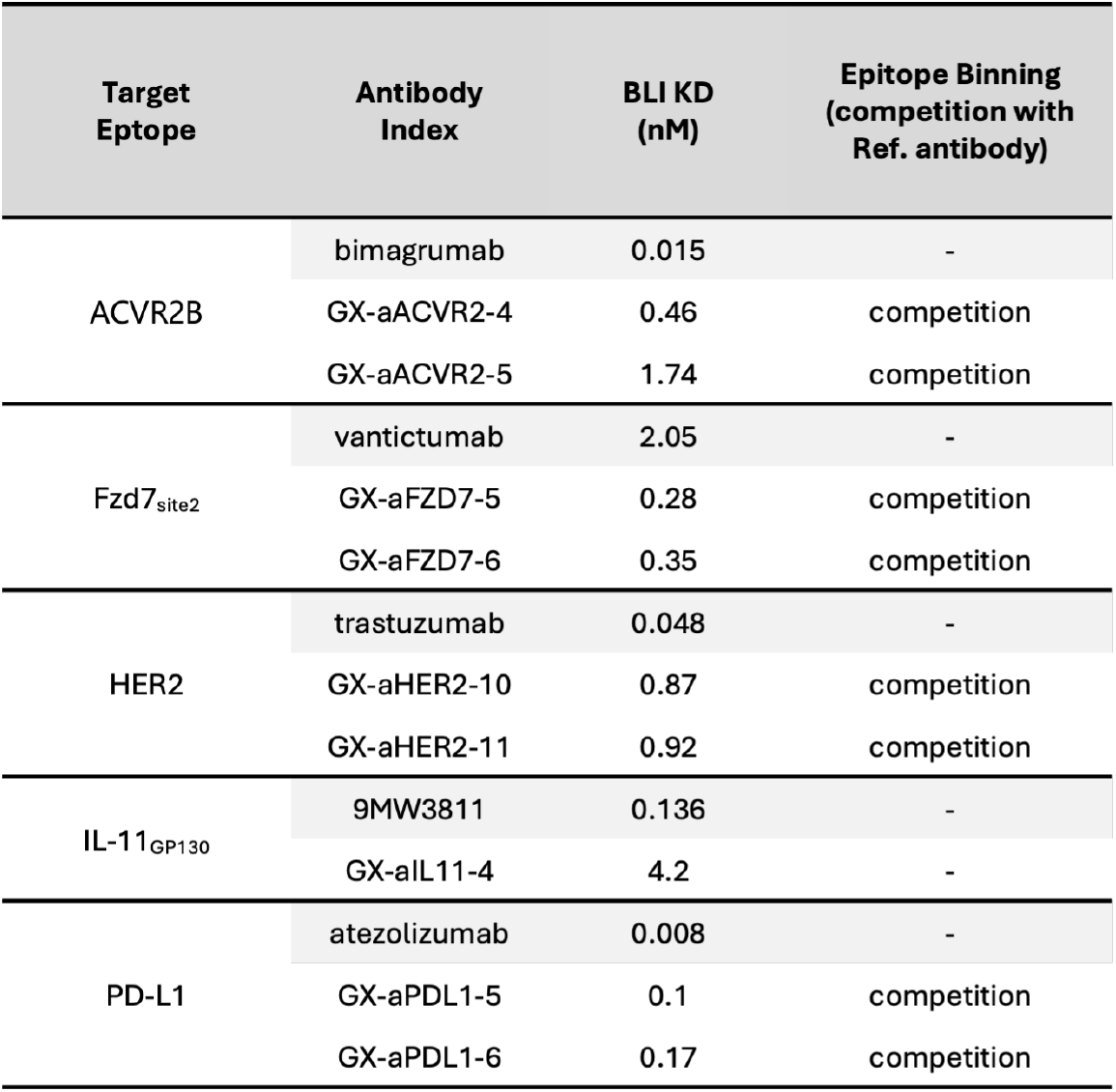
*In Vitro* Binding Characterization Profiles of Designed Antibodies in the IgG Format.

| Target Epitope | Antibody Index | BLI KD (nM) | Epitope Binning (competition with Ref. antibody) |
| --- | --- | --- | --- |
| ACVR2B | bimagrumab | 0.015 | - |
|  | GX-aACVR2-4 | 0.46 | competition |
|  | GX-aACVR2-5 | 1.74 | competition |
| Fzd7<sub>site2</sub> | vantictumab | 2.05 | - |
|  | GX-aFZD7-5 | 0.28 | competition |
|  | GX-aFZD7-6 | 0.35 | competition |
| HER2 | trastuzumab | 0.048 | - |
|  | GX-aHER2-10 | 0.87 | competition |
|  | GX-aHER2-11 | 0.92 | competition |
| IL-11<sub>GP130</sub> | 9MW3811 | 0.136 | - |
|  | GX-aIL11-4 | 4.2 | - |
| PD-L1 | atezolizumab | 0.008 | - |
|  | GX-aPDL1-5 | 0.1 | competition |
|  | GX-aPDL1-6 | 0.17 | competition |

To visually illustrate the distinct properties and therapeutic potential of these candidates, Figures 4 and 5 present a characterization overview for two representative binders targeting Fzd7 and PD-L1. Comprehensive data for representative binders from the other successful epitopes are provided in Supplementary Figure S9.

**Figure 4.**
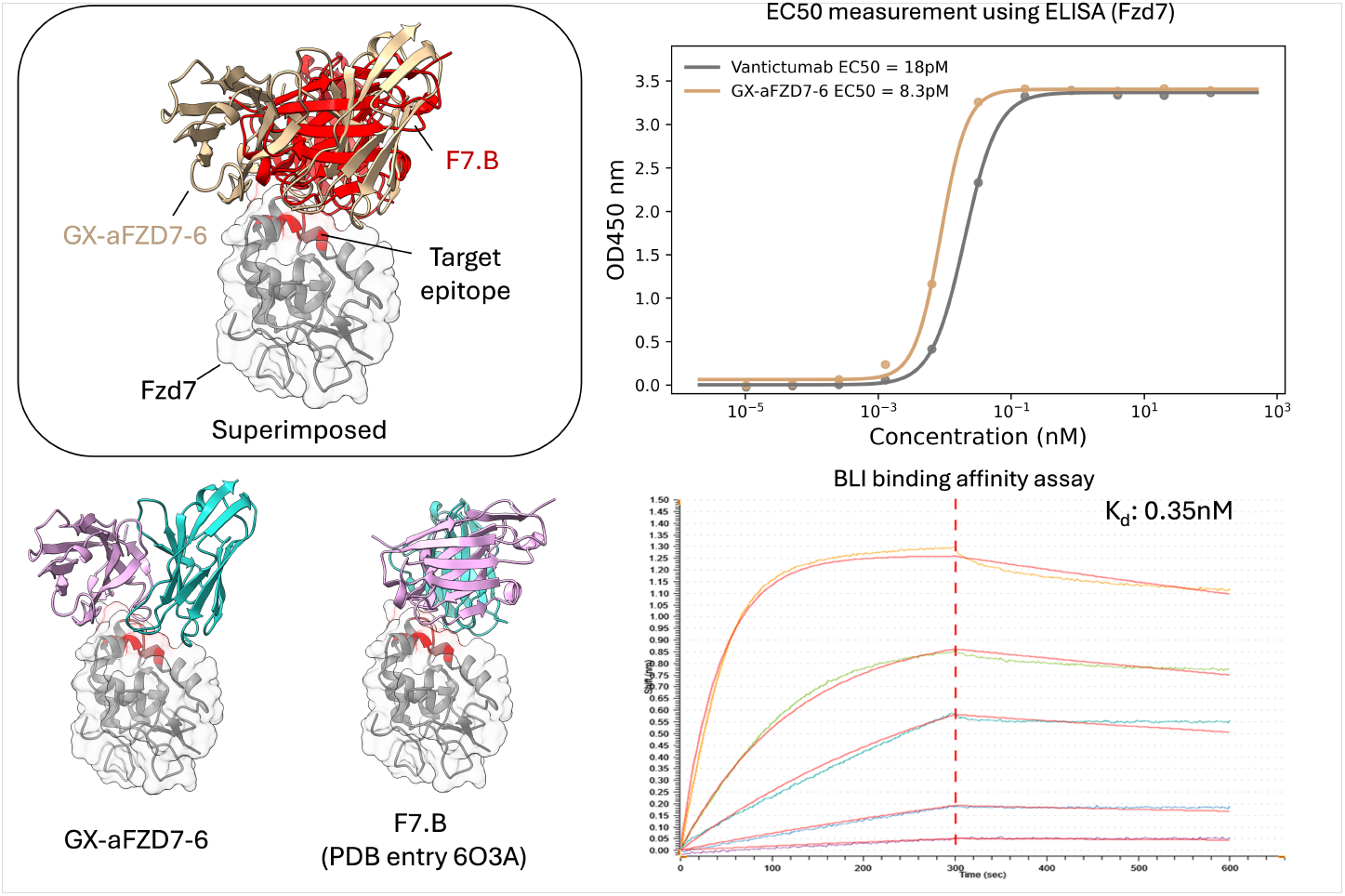
Characterization of a Representative Fzd7 High-Affinity Binder (GX-aFZD7-6). Structural comparison features the predicted binding orientation of GX-aFZD7-6 overlaid with the most structurally similar PDB complex (PDB ID: 6O3A), with a framework C-*α* RMSD of 29.7 Å. Binding affinity is quantified by ELISA (EC_50_ = 8.3 pM) and BLI-based kinetics (K_d_ = 0.35 nM). This binder demonstrates sequence novelty, exhibiting 52.8% max CDR and 50.0% max CDR H3 sequence identity relative to SAbDab entries.

**Figure 5.**
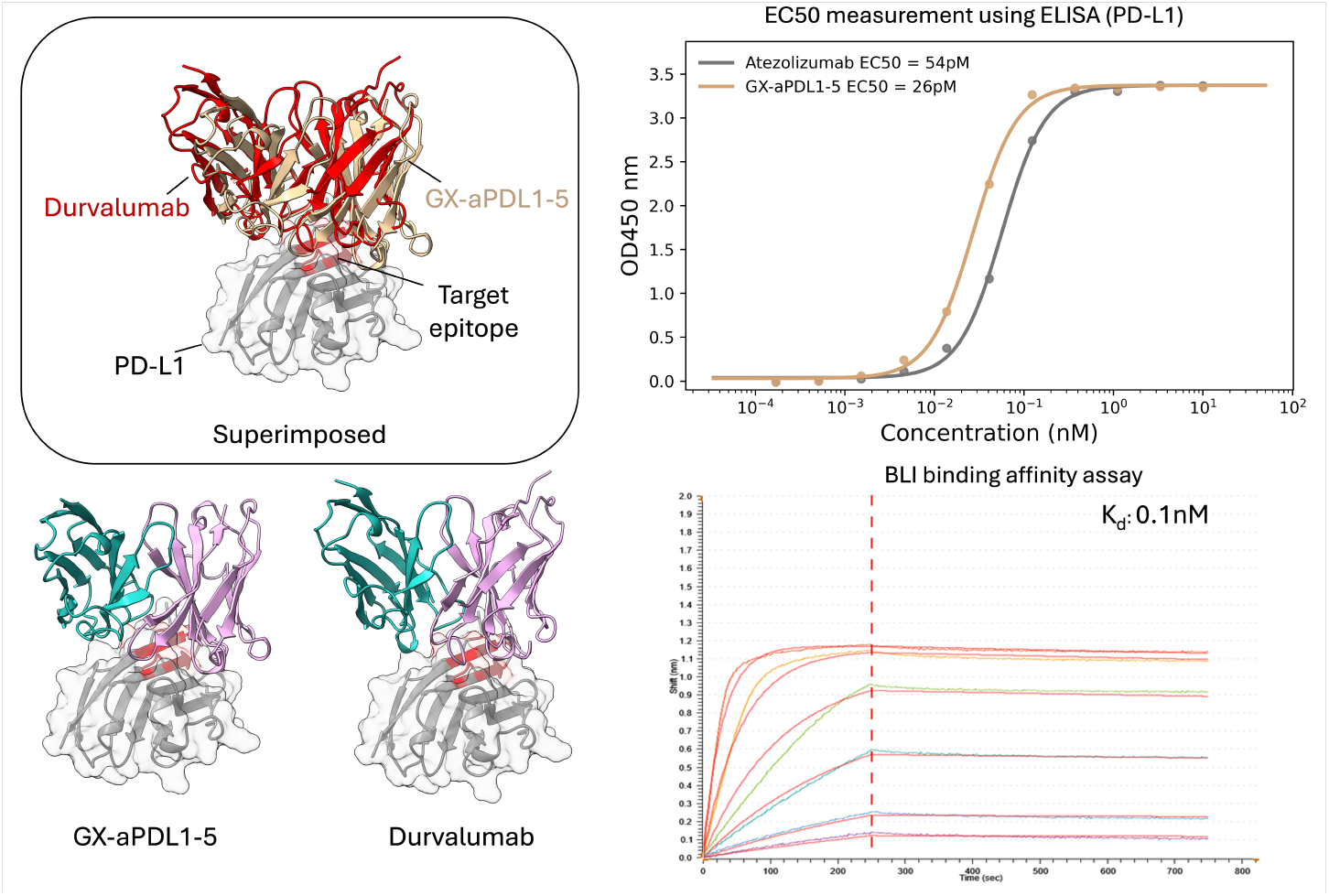
Characterization of a Representative PD-L1 High-Affinity Binder (GX-aPDL1-5). Structural comparison features the predicted binding orientation of GX-aPDL1-5 overlaid with the most structurally similar PDB complex (PDB ID: 5X8M), with a framework C-*α* RMSD of 7.9 Å. Binding affinity is quantified by ELISA (EC_50_ = 26 pM) and BLI-based kinetics (K_d_ = 0.1 nM). This binder demonstrates sequence novelty, exhibiting 57.6% max CDR and 55.6% max CDR H3 sequence identity relative to SAbDab entries.

Epitope binning via biolayer interferometry (BLI) confirmed that selected binders for PD-L1, HER2, ACVR2B, and Fzd7 all successfully competed with reference antibodies known to bind the same target region. (A suitable reference antibody for epitope binning was not available for IL-11.) This validated that the *in silico* design process accurately directed the antibodies to the designated epitope or a closely overlapping region for all four targets tested (Table 1).

### 2.5 Atomic-Level Structural Analysis of Designed Interactions

Analysis of the predicted complex structures for the binders provides key insights into the structural basis of their binding. Figure 6 illustrates the predicted interactions for two representative high-affinity binders, one targeting Fzd7 and one targeting PD-L1. The anti-Fzd7 candidate (GX-aFZD7-6) is predicted to achieve its affinity through a specific combination of hydrogen bonds and hydrophobic interactions that precisely engage key residues within the targeted epitope. In the case of GX-aFZD7-6, the interaction with Fzd7 includes a salt bridge between an Asp residue in the CDR-H1 loop and K96 of Fzd7, while the CDR-H2 and CDR-H3 loops contribute a cluster of hydrophobic contacts around F97 that help stabilize the binding interface (Figure 6A). Similarly, the anti-PD-L1 candidate (GX-aPDL1-5) is predicted to form stable salt-bridge contacts together with a broad network of hydrophobic interactions at the binding interface. PD-L1 recognition involves hydrogen bonds between two Thr residues in the CDR-H3 loop and a charged patch centered on D61, together with a salt bridge formed by an Arg residue in the CDR-H3 loop and R125, supported by additional hydrophobic interactions across the interface (Figure 6B).

**Figure 6.**
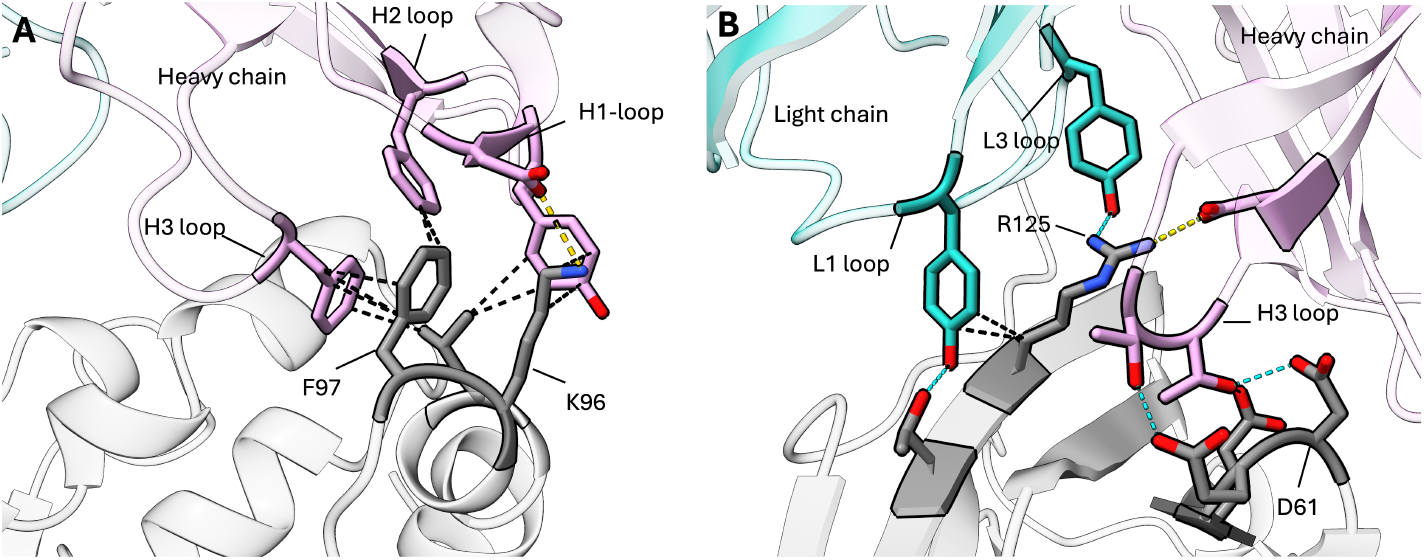
Interaction Analysis of Designed Antibody Binders. (A) Key interactions formed by the anti-Fzd7 antibody (GX-aFZD7-6). The binding pose features a salt bridge between an Asp residue (CDR-H1) and K96 of Fzd7, supplemented by hydrophobic contacts mediated by the CDR-H2 and CDR-H3 loops surrounding F97. (B) Key interactions formed by the anti-PD-L1 antibody (GX-aPDL1-5). Interactions include hydrogen bonding between two Thr residues (CDR-H3) and a charged patch centered on D61 of PD-L1. Additionally, a salt bridge occurs between an Arg residue (CDR-H3) and R125, alongside distributed hydrophobic interactions across the binding interface.

### 2.6 Cross-Platform Benchmark

Having established the Precision-Scale Workflow on therapeutically relevant epitopes, we next evaluated an independent set of common target proteins selected for comparison with the Chai-2 [5] and JAM-2 [8] preprints. This cross-platform benchmark set comprised nine target proteins: alpha hemoglobin stabilizing protein (AHSP), prolactin (PRL), low affinity immunoglobulin gamma Fc region receptor III-B (FCG3B), endothelial protein C receptor (EPCR), parvin alpha (PARVA), mitogen-activated protein kinase 8 (MK08), ring finger protein 43 (RNF43), CD226 molecule (CD226), and interleukin 3 (IL3).

For each target protein, the top 50 GaluxDesign candidates were synthesized and tested directly as full-length IgGs by 100 nM single-point ELISA. Using the same criteria as Subsection 2.2, candidates were classified as binders(relaxed) or binders(strict). Across the nine benchmark targets, GaluxDesign identified 79 binders(relaxed) and 49 binders(strict), corresponding to normalized hit rates of 17.6% and 10.9%, respectively. At the target level, GaluxDesign yielded binders(strict) for eight of the nine targets, with IL3 as the only target for which no binder was obtained in the present campaign. The full on-target/off-target ELISA distributions for these experiments are provided in Supplementary Figure S8.

BLI-based kinetic analysis further confirmed apparent binding affinities in the low-to-mid nanomolar range for seven of the nine benchmark targets. Representative apparent K_d_ values and normalized BVP scores for the selected binders are summarized in Table 2, and the corresponding fitted BLI profiles are provided in Supplementary Figure S10. Notably, most normalized BVP scores were close to 0, consistent with trastuzumab-like polyspecificity profiles and indicating no substantial nonspecific binding tendency among the characterized binders. The full-length heavy- and light-chain amino-acid sequences of the representative AHSP and PARVA binders are provided in Supplementary Table S3. Sequences from the other six successful targets are not disclosed here because of their potential therapeutic relevance.

**Table 2.** BLI binding characterization and normalized BVP scores of representative binders from the cross-platform benchmark.

| Target | Antibody Index | BLI $K_d$ (nM) | Normalized BVP score<sup>†</sup> |
| --- | --- | --- | --- |
| PRL | GX-aPRL-1 | 1.5 | 0.100 |
| PARVA | GX-aPARVA-1 | 1.9 | -0.022 |
| FCG3B | GX-aFCG3B-1 | 2.7 | 0.094 |
| AHSP | GX-aAHSP-1 | 4.7 | 0.284 |
| RNF43 | GX-aRNF43-1 | 29.6 | n.d. |
| EPCR | GX-aEPCR-1 | 37.0 | -0.007 |
| CD226 | GX-aCD226-1 | 59.4 | -0.001 |
| MK08 | GX-aMK08-1 | n.q. | 0.064 |
| - | visilizumab | - | 1.000 |
| - | trastuzumab | - | 0.000 |

<sup>†</sup>Normalized BVP scores were calculated from replicate-averaged OD<sub>450</sub> values after BVP-only blank subtraction, followed by linear normalization using trastuzumab = 0.0 and visilizumab = 1.0. Negative values indicate BVP signals below the trastuzumab control. n.d., not determined. n.q., not quantifiable.

Overall, these results support the competitive target-level performance of the Precision-Scale Workflow in this cross-platform benchmark setting. GaluxDesign identified experimentally validated antibody binders for eight of the nine benchmark targets (89%), compared with five of nine targets for JAM-2 (56%) and four of nine targets for Chai-2 (44%) (Figure 7). Although antibody format and assay configuration differ across the reported results, this target-level comparison supports the favorable performance of GaluxDesign under a fixed 50-candidate budget per target.

**Figure 7.**
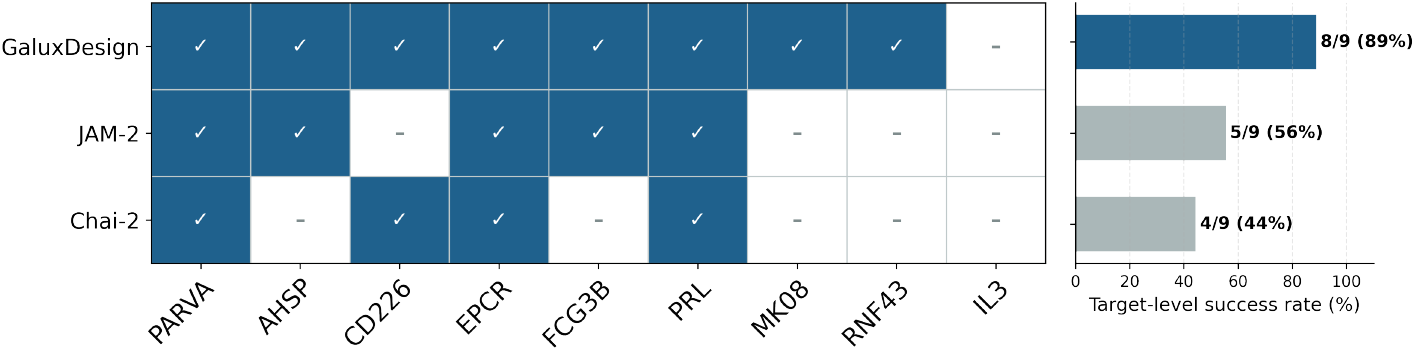
*De novo* antibody design across shared benchmark targets compared with reported Chai-2 and JAM-2 results. Check marks indicate that at least one experimentally validated binder was reported for the corresponding target, whereas dashes indicate that no binders were identified. GaluxDesign and JAM-2 results are based on full-length antibodies, whereas Chai-2 results are based on scFv designs.

## 3 Conclusion

This study demonstrates that GaluxDesign can identify high-affinity full-length IgG binders from a minimal experimental set using the Precision-Scale Workflow. In the therapeutic epitope set, testing only 50 candidates per epitope yielded a 31.5% binders(relaxed) rate and a 10.5% binders(strict) rate, with picomolar-to-nanomolar affinity binders identified for seven of eight epitopes.

We further evaluated the same workflow on nine shared benchmark targets selected for comparison with the Chai-2 and JAM-2 preprints. At the target level, GaluxDesign identified binders for eight of nine targets, including BLI-characterized binders with apparent affinities in the low-to-mid nanomolar range for seven targets, supporting its competitive performance in this external benchmark setting.

Together, these findings demonstrate the breadth and efficiency of GaluxDesign at the Precision Scale. Under a fixed 50-candidate budget, the platform can progress from *in silico* design to experimentally validated, target-specific full-length IgG binders across distinct targets, supporting rapid and resource-efficient antibody lead generation.

## 4 Methods

### 4.1 Computational design and candidate selection

*De novo* antibody variable region (Fv) sequences were computationally generated using the GaluxDesign platform [1]. For the therapeutic epitope set (Subsection 2.1 to 2.4), candidate binding sites were defined by manually selecting two to six residues for each target protein based on functional relevance and structural accessibility. For the cross-platform benchmark set (Subsection 2.6), candidate binding sites were selected using the same criteria, incorporating information from known natural binding interfaces or complex structures when available. For each specified site, GaluxDesign generated a diverse pool of candidates, from which the top 50 unique sequences were prioritized based on *in silico* scoring functions for experimental validation.

### 4.2 IgG production and purification

For the initial screening, *de novo* designed antibodies were produced externally by Biointron (China) using the HEK293 expression system. Subsequently, for those antibodies selected for further binding analyses, human codon-optimized gBlock™ (IDT) coding for the heavy chain (HC) including the VH and CH regions, and light chain (LC) including the VL and CL regions was first inserted into pcDNA™3.4-based plasmid (X11 for HC and X15 for LC) using restriction enzyme cloning. The plasmid DNA (0.5 μg) for each HC and LC dissolved in 40 μL of sterile water was then transfected into cultured Expi293F cells with ExpiFectamine™ reagent and cultured for 4 days at 37 °C, 8% CO2, 125 rpm and 80% humidity to make IgG. The culture supernatant was centrifugated and filtered through a 0.22 μm filter. The titer of recombinant IgG was analyzed using BLI. The binder antibodies were subsequently purified using MabSelect™ (Cytiva) affinity chromatography resin. Antibody concentration was determined by measuring the absorbance at A280 using Nanodrop (Thermo Fisher Scientific), and the molecular weight of the antibody was confirmed on SDS-PAGE analysis.

### 4.3 Screening by ELISA and binder criteria

A single-point (100 nM concentration) ELISA was performed for binder screening across both target sets evaluated in this report: the therapeutic epitope set and the cross-platform benchmark set. For each epitope or target protein, the top 50 GaluxDesign candidates were synthesized as full-length IgGs and tested at 100 nM antibody concentration.

For the therapeutic epitope set, the optimized antigen was coated on a 96-well plate and incubated for 12–16 hours. After washing with PBS-T buffer (1 × DPBS, 0.05% Tween20), the plate was blocked with 3% skim milk in PBS-T buffer for one hour. The antibody at 100 nM was then added to the antigen-coated plate for two hours. A secondary antibody (anti-human IgG antibody-HRP conjugation) was added for one hour. The signal was developed by adding TMB substrate for 30 minutes, followed by addition of a stop solution to terminate the reaction. The absorbance was measured at OD_450_.

For the cross-platform benchmark set, the same single-point ELISA workflow was applied with target-optimized antigen coating conditions. On-target signals were corrected by blank subtraction, and specificity was evaluated using an off-target control antigen under the same assay conditions.

The logic for binder classification was as follows:

- binders(relaxed): Defined as designs meeting
  1. 100 nM on-target signal (ELISA OD) *>* 0.2 AND
  2. “off-target” criteria (relaxed) (2-1) 100 nM off-target signal was *<* 0.2 OR (2-2) the off-target signal was *<* 10% of the on-target signal.

- binders(strict): Defined as designs meeting
  1. 100 nM on-target signal *>* 1.8 (50% of OD_max_), AND
  2. off-target signal *<* 10% of on-target signal.

### 4.4 Novelty and diversity analysis

Sequence novelty and diversity were calculated based on the CDR sequences (Chothia definition, CDRs 1-3) of the heavy and light chains. Sequence identity was computed using Needle [10]. Structural orientation novelty was assessed by first aligning the target protein of the designed complex to the antigen component of relevant antibody-antigen complexes in the PDB. Based on this alignment, the Framework (non-CDR) C-*α* RMSD was then calculated between the designed antibody and reference antibodies.

### 4.5 Binding affinity (EC_50_) analysis using ELISA

To determine binding affinity, an ELISA was performed for EC_50_ estimation. The optimized antigen was coated on a 96-well plate and incubated overnight at 4 °C. After incubation, the plate was washed with PBS-T buffer (1 × DPBS, 0.05% Tween20).

The wells were then blocked with 3% skim milk in PBS-T buffer for 1 hour at room temperature (RT). A serial dilution of the test antibody was added to the antigen-coated wells and incubated for 2 hours at RT. Following another wash step, an HRP-conjugated anti-human IgG secondary antibody was added and incubated for 1 hour at RT. The signal was developed by adding TMB substrate and allowing the reaction to proceed for 30 minutes, after which a stop solution was added to terminate the reaction. Absorbance was measured at OD_450_ using a microplate reader.

### 4.6 Binding affinity analysis using BLI

All binding assays were carried out on a Gator Prime instrument (GatorBio) using 1 × PBST (1 × PBS, 0.05% Tween-20) as the running and wash buffer. Sensors were pre-equilibrated in buffer prior to use, and washing was performed between each assay step. The assay was conducted under optimized conditions for each target to ensure sensitivity and reproducibility.

#### ACVR2B

A His-tagged ACVR2B was immobilized on Ni-NTA sensors at 2.5 μg/mL. Association was measured by exposing the loaded sensors to the anti-ACVR2B antibody at concentrations starting from 5 μg/mL in a two-fold serial dilution series for 300 s, followed by dissociation in 1*×*PBST for 300 s.

#### Fzd7

A His-tagged Fzd7 was immobilized on Ni-NTA sensors at 5 μg/mL. Association was measured by exposing the loaded sensors to the anti-Fzd7 antibody at concentrations starting from 7.5 μg/mL in a three-fold serial dilution series for 300 s, followed by dissociation in 1*×*PBST for 300 s.

#### HER2

A biotinylated HER2 was immobilized on streptavidin (SA) sensors at 1.25 μg/mL. Association was measured by exposing the loaded sensors to anti-HER2 antibody at concentrations starting from 7.5 μg/mL in a 2-fold serial dilution series for 200 s, followed by dissociation in 1*×*PBST for 600 s.

#### IL-11

The anti-IL-11 antibody was captured at 2.5 μg/mL on an HFc probe, with association measured with IL-11 starting at 2 μg/mL in a three-fold serial dilution series for 500 s, followed by dissociation in 1*×*PBST for 500 s.

#### PD-L1

A His-tagged PD-L1 was immobilized on Ni-NTA sensors at 6.5 μg/mL. Association was measured by exposing the loaded sensors to the anti-PD-L1 antibody at concentrations starting from 7.5 μg/mL in a 2-fold serial dilution series for 250 s, followed by dissociation in 1*×*PBST for 500 s.

Raw sensorgrams from all experiments were reference-subtracted and further double-referenced using either a reference channel or the lowest-concentration well of the same probe type. Kinetic parameters were obtained by global fitting to a 1:1 binding model, in which both association and dissociation phases were simultaneously fitted to derive on-rate constant (k_on_), off-rate constant (k_off_), and the equilibrium dissociation constant (K_d_=k_off_/k_on_). Normalization of loading responses was applied prior to fitting to minimize sensor-to-sensor variability. Data analysis was performed using GatorBio analysis software to determine binding characteristics.

Selected ELISA-positive antibodies from the cross-platform benchmark were characterized using the same BLI analysis framework on an OCTET RH96 system (Sartorius). Unless otherwise specified, antibodies were captured on AHC2 biosensors at 8.3 nM, followed by association with the corresponding antigen. Target proteins were tested in a two-fold serial dilution series for 500 s, followed by dissociation in 1 × kinetics buffer for 500 s. Top analyte concentrations were 100 nM for most proteins, except RNF-43 (1 μM) and AHSP-1 (50 nM). Apparent K_d_ values were reported only when the fitted BLI profiles supported quantitative kinetic interpretation.

### 4.7 Epitope binning by BLI

Epitope binding analysis was performed using BLI on a Gator Prime system (GatorBio). This assay determined whether the designed antibodies bind to the same epitope as a reference antibody, as well as to assess their binding affinity. A general competitive binding assay was performed where the antigen was immobilized first. Following baseline stabilization, the first antibody was loaded onto the antigen-coated biosensors. A subsequent association step with a second antibody was then performed to evaluate simultaneous binding. The absence of a binding response from the second antibody indicated that the two antibodies recognize overlapping epitopes or sterically hinder each other. A detectable binding response indicated they bind to distinct epitopes.

### ACVR2B

ACVR2B was immobilized on Ni-NTA biosensors at 6.5 μg/mL. Following baseline stabilization in 1 × PBST, the primary antibody (reference bimagrumab or an engineered variant) was associated with the ACVR2B-coated sensors at 15 μg/mL for 900 s. A second association step was then performed by introducing the alternate antibody at the same concentration for an additional 900 s to determine whether both antibodies.

### Fzd7

Fzd7 was immobilized on Ni-NTA biosensors at 5 μg/mL. Following baseline stabilization in 1 × PBST, the primary antibody (reference F7.B (PDB entry 6O3A) or an engineered variant) was associated with the Fzd7-coated sensors at 15 μg/mL for 900 s. A second association step was then performed by introducing the alternate antibody at the same concentration for an additional 900 s.

#### HER2

Biotinylated HER2 was immobilized on streptavidin (SA) biosensors at 5 μg/mL. Following baseline stabilization in 1 × PBST, the primary antibody (reference trastuzumab or an engineered variant) was associated with the HER2-coated sensors at 15 μg/mL for 900 s. A second association step was then performed by introducing the alternate antibody at the same concentration for an additional 900 s.

#### PD-L1

PD-L1 was immobilized on Ni-NTA biosensors at 6.5 μg/mL. Following baseline stabilization in 1 × DPBS, the primary antibody (reference atezolizumab or an engineered variant) was associated with the PD-L1-coated sensors at 15 μg/mL for 900 s. A second association step was then performed by introducing the alternate antibody at the same concentration for an additional 900 s to assess whether antibodies could bind simultaneously.

### 4.8 Poly-specificity assay

Baculovirus particle (BVP) ELISA was used as an orthogonal polyspecificity-related candidate-quality assay for the representative binders summarized in Table 2. For each antibody, replicate OD_450_ values were averaged. BVP-only wells were used as blank controls and subtracted from the sample, trastuzumab, and visilizumab signals within the same assay block. The blank-corrected values were then linearly normalized such that trastuzumab corresponded to 0.0 and visilizumab corresponded to 1.0, yielding the Normalized BVP score reported in Table 2:

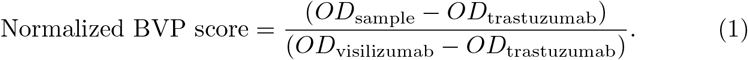

Note that the scores below 0 indicate BVP signals lower than trastuzumab under the same assay conditions.

## 5 Author Contributions

T. P., C. S., J. W., and J. Y. conceived the study and coordinated the collaborations. J. G., S. W. H., Sohee K., C. L., Danyeong L., Dohoon L., J. L., J. N., S. R., J. S., J. W., H. W., and J. Y. developed the predictive and generative AI methods for *de novo* protein design. H. C., K. C., D. G., M. H., Sehan K., Suyeon K., D. K. L., S. O., E. P., S. P., E. R., D. H. S., and M. Y. S. conducted the wet-lab validations. K. C., M. H., Dohoon L., C. S., J. W., H. W., and J. Y. wrote the initial draft of the manuscript. All authors contributed to the final version.

## 6 Competing Interests Statement

The authors are executives or employees of Galux Inc. and may have financial interests related to this research.

---

**Figure S8.**
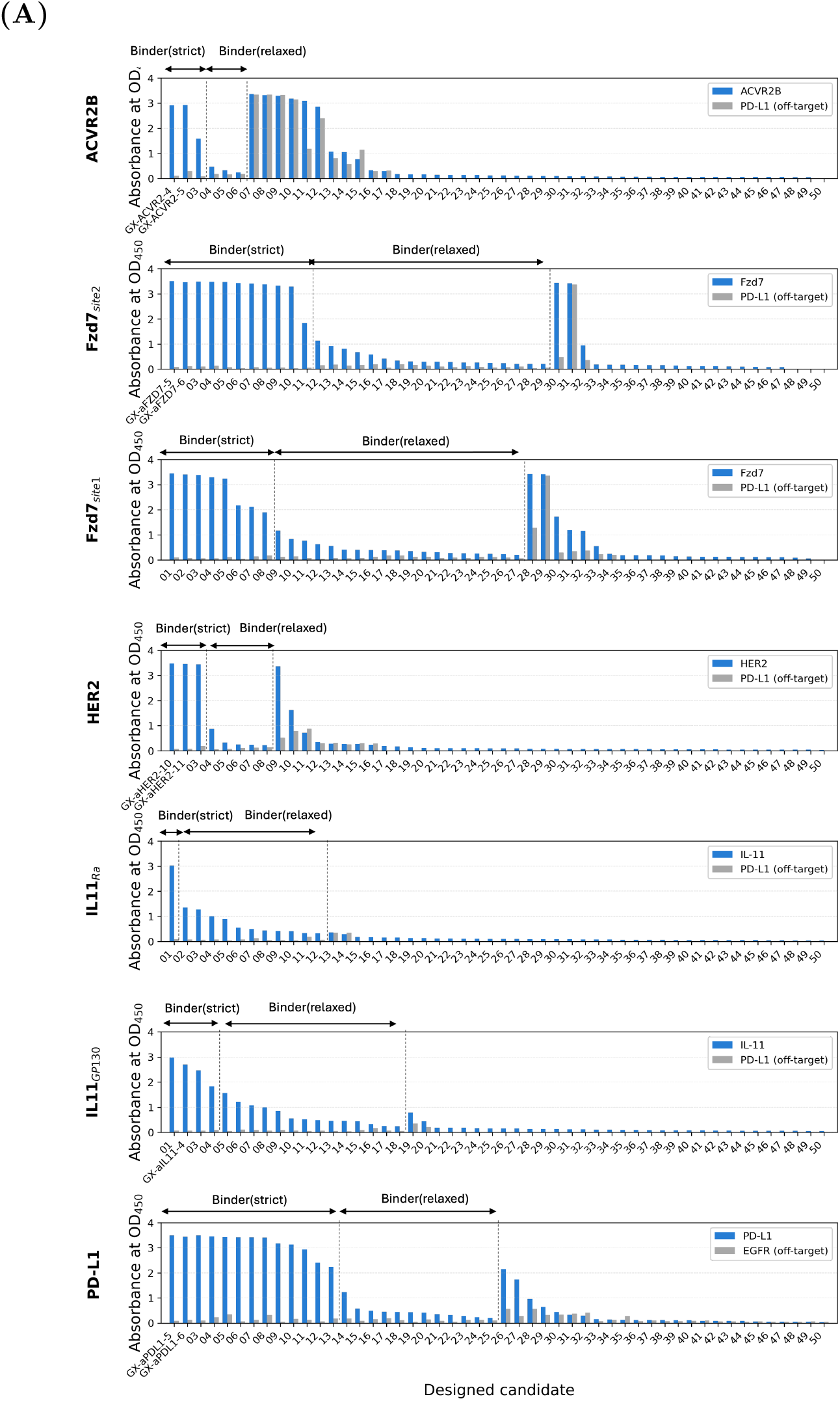
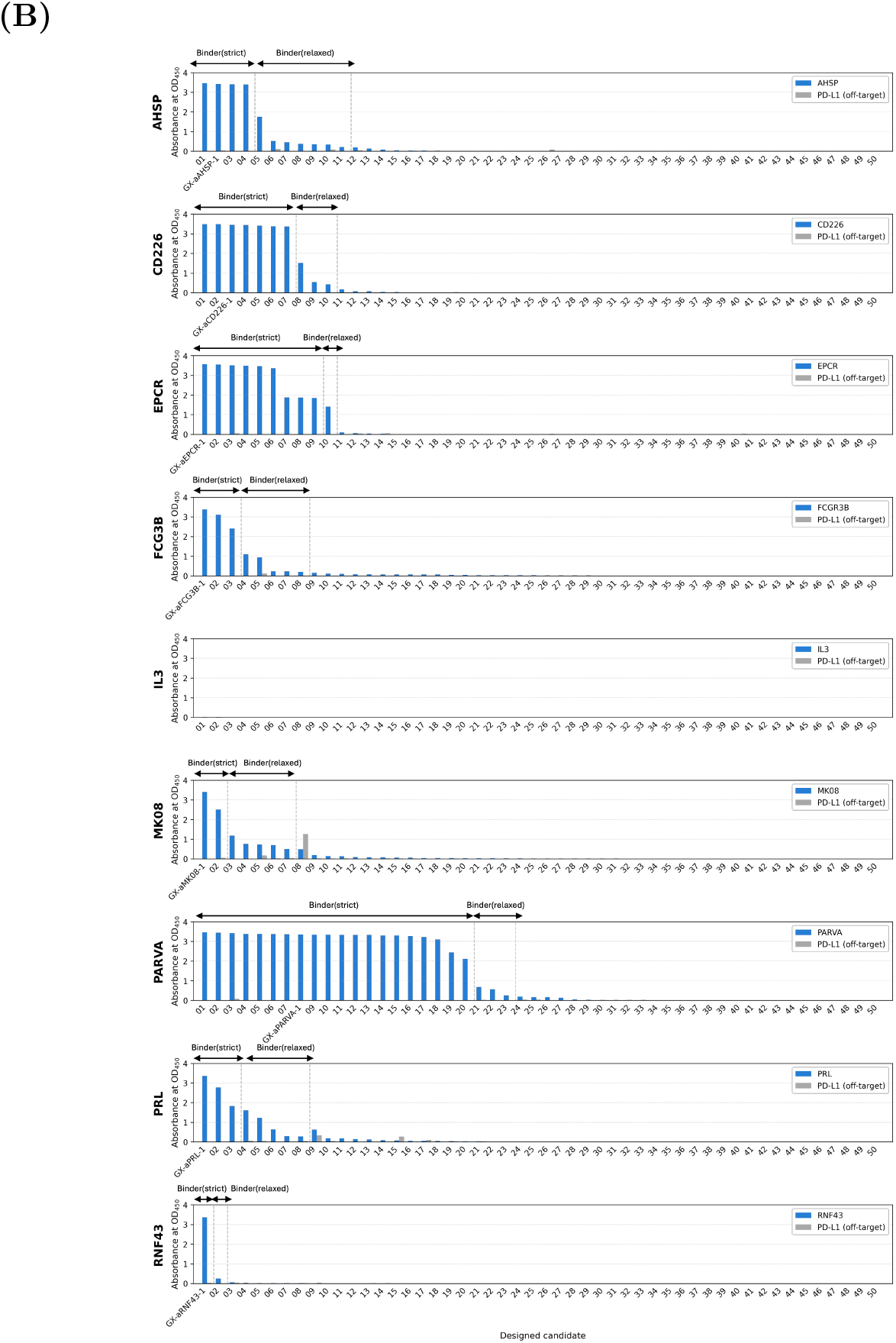
On-target (OD_450_) versus off-target signals from the 100 nM single-point ELISA screening. (A) Screening results for the therapeutic epitope set. Each point represents a single design from one of the eight target epitopes, plotted by its on-target signal (blue) and off-target signal (gray). (B) Screening results for the cross-platform benchmark target set. Each point represents a single design from one of the benchmark target proteins, plotted using the same on-target and off-target signal definitions. The same ELISA criteria were applied to classify binders(relaxed) and binders(strict).

**Figure S9.**
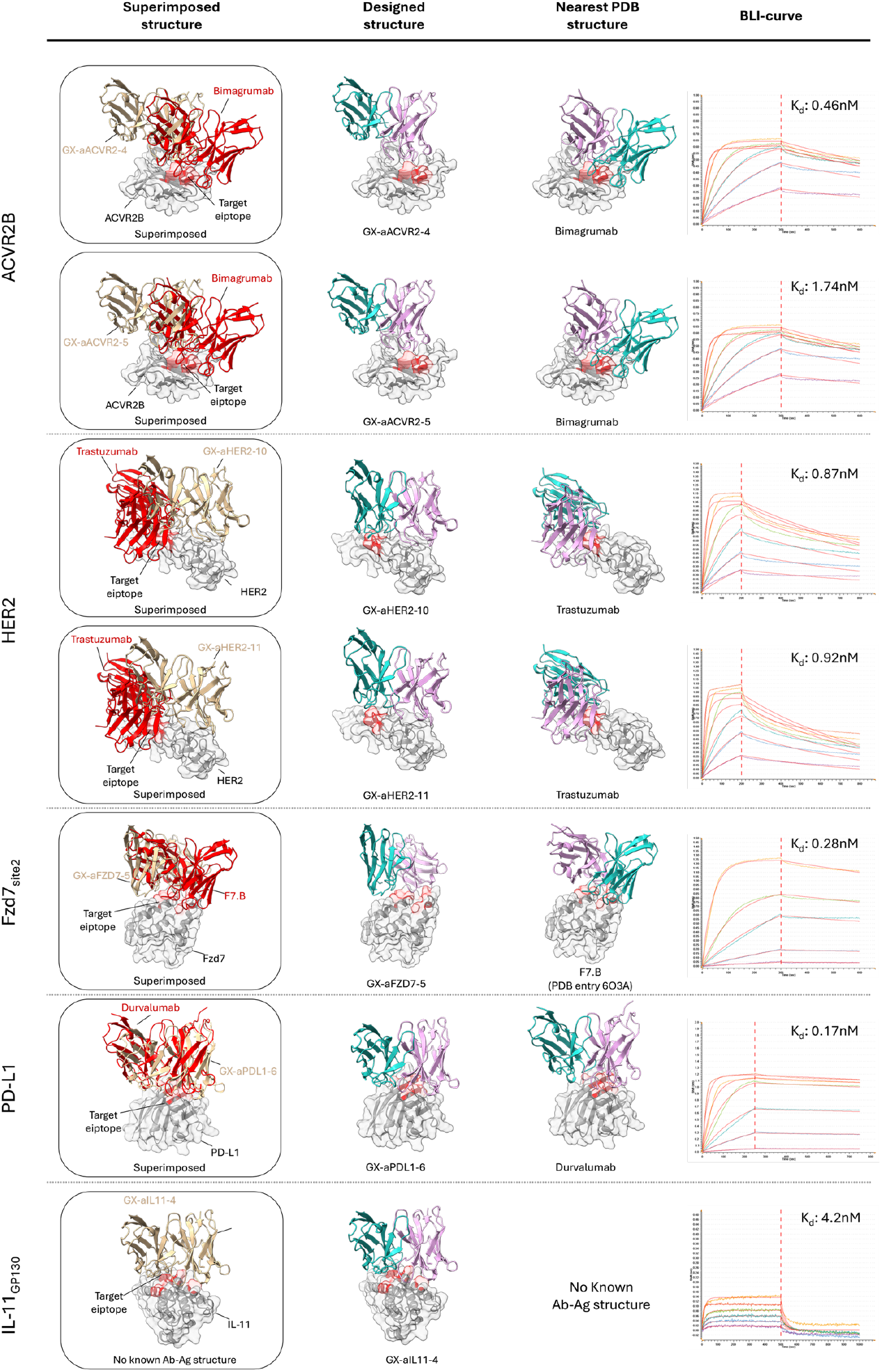
Structural novelty and binding kinetics for each *in silico* designed antibody. (Left) Superimposition of the computationally designed antibody– target complex (in tan) and the known antibody–target complex with the closest binding pose (in red) confirming novelty of the binding pose. (Right) Biolayer Interferometry (BLI) sensorgrams and the calculated dissociation constant (K_d_) value (*R*^2^ *>* 0.95) demonstrating the sub-to single-digit nanomolar binding kinetics of the designed antibody.

**Figure S10.**
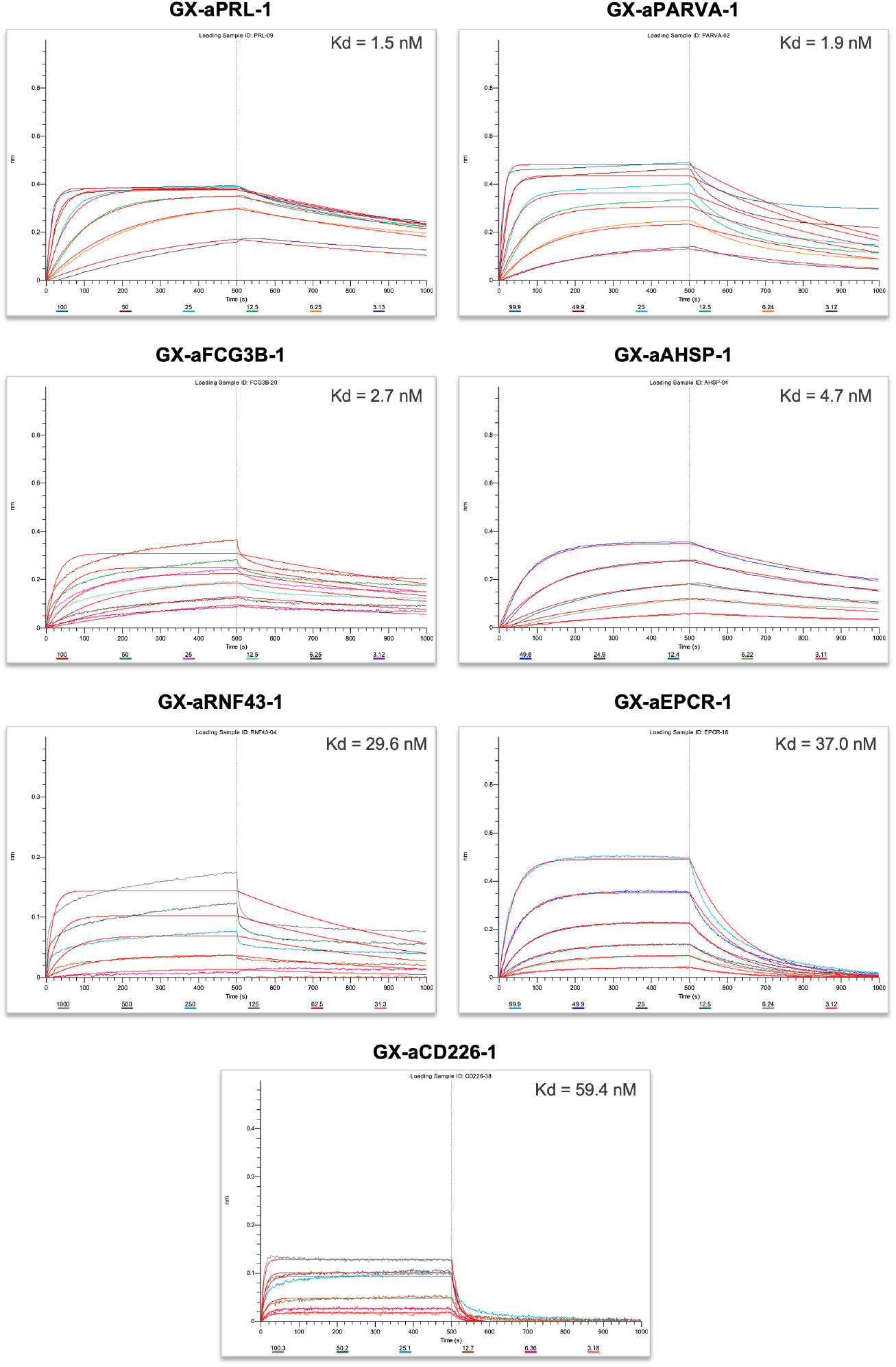
Fitted BLI profiles (*R*^2^ *>* 0.95, except GX-aRNF43-1, *R*^2^ = 0.94) for representative binders from the cross-platform benchmark. Profiles are shown for the antibody candidates summarized in Table 2.

**Table S3.**
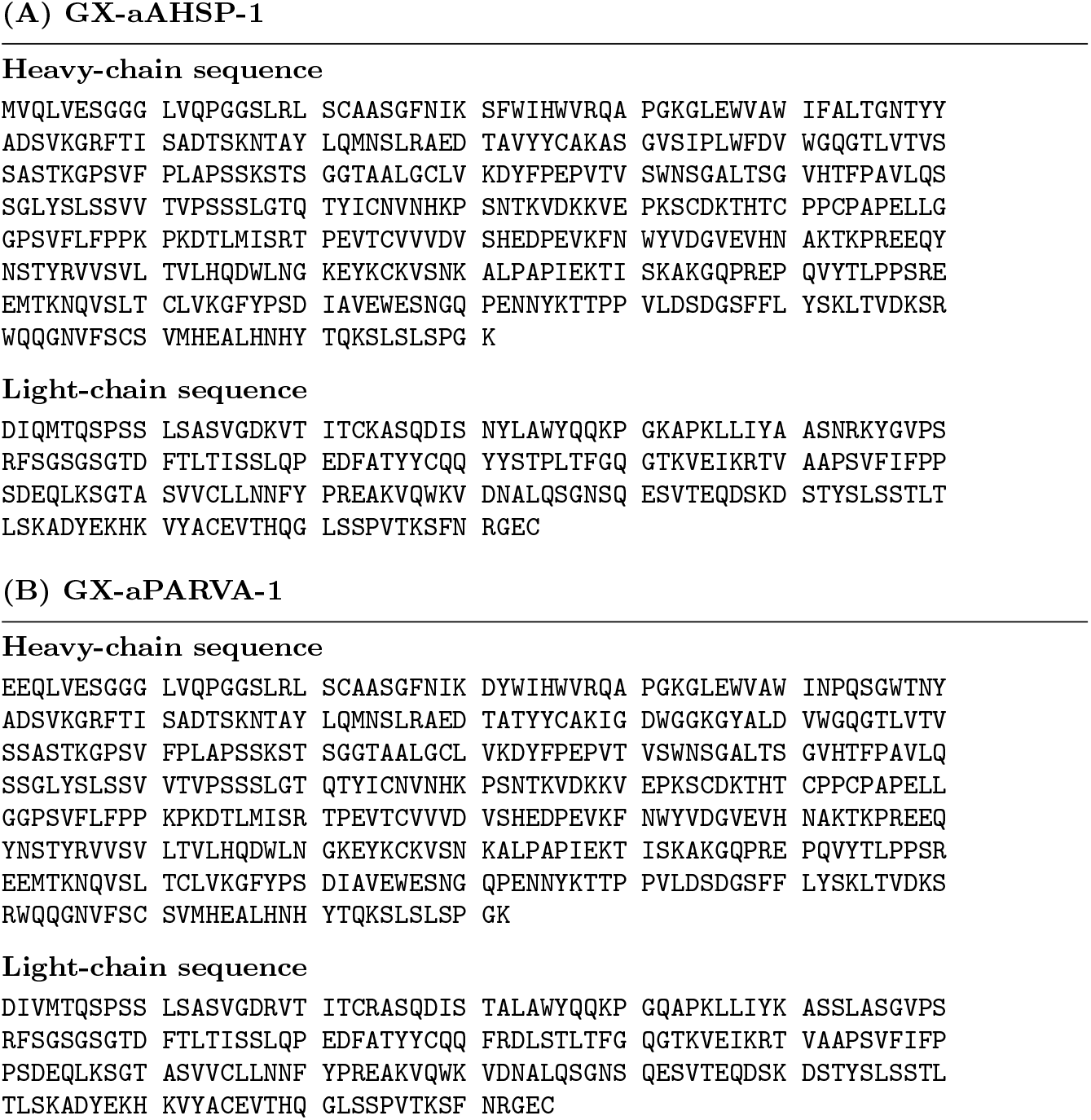
Full-length sequences of two representative binders from the cross-platform benchmark.

## Notes

### Summary of Updates

This revised manuscript includes a major update to Section 2.6 with the addition of a cross platform comparison study using nine shared antibody design targets selected from publicly disclosed studies. We experimentally evaluated 50 full length IgG candidates per target and compared target level binder discovery outcomes with publicly reported results from recent de novo antibody design studies. The revised manuscript now includes new ELISA screening results, BLI based kinetic characterization, normalized BVP analysis, additional supplementary figures and tables, and expanded methodological descriptions related to the cross platform evaluation workflow. The Results and Discussion and Conclusion sections were updated accordingly to reflect the expanded validation scope and comparative findings.

